# Levetiracetam inhibits SV2A-synaptotagmin interaction at synapses that lack SV2B

**DOI:** 10.1101/640185

**Authors:** Kristine Ciruelas, Daniele Marcotulli, Jane M Sullivan, Sandra M Bajjalieh

## Abstract

Epilepsy remains a difficult-to-treat neurological disorder prompting the need for new therapies that work via alternate mechanisms. Levetiracetam (**LEV**) is the first in a series of anti-epilepsy drugs that target presynaptic functioning. LEV binds the synaptic vesicle protein SV2A, and has been shown to decrease neurotransmitter release in hippocampal slices. The molecular basis of LEV action is unknown, however, and direct effects of LEV on SV2A function remain to be determined. SV2A is the most widely expressed paralog of a three-gene family (SV2A, B, C) that is variably co-expressed throughout the CNS. All three SV2s bind the calcium sensor protein synaptotagmin and SV2 plays a crucial role in synaptotagmin stability and trafficking. Here we addressed the action of LEV at the cellular and molecular level asking whether the presence of non-LEV binding SV2 paralogs influences drug action and whether LEV impacts SV2A’s role in synaptotagmin function. We report that LEV altered short-term synaptic plasticity in isolated neurons from SV2B knockout but not wild-type mice, mimicking the loss of SV2 function. Similarly, LEV reduced SV2A binding to synaptotagmin only in the absence of SV2B. Furthermore, LEV reduced and slowed the internalization of synaptotagmin in neurons cultured from SV2B KO but not WT mice. Taken together, these findings suggest that LEV alters synaptic release probability by disrupting SV2’s regulation of synaptotagmin selectively in neurons that express only SV2A. Neurons that meet this requirement include most inhibitory neurons and the granule cells of the dentate gyrus, two classes of neuron implicated in epilepsy.

## Introduction

Levetiracetam (Keppra™, **LEV**) is a second-generation anti-epileptic drug (**AED**) that lacks many of the adverse side effects of classic AEDs. In addition to its use for epilepsy, LEV is being used to treat a range of neurological disorders and has recently been found to be a promising treatment for cognitive impairment (1). Hindering its further development is the fact that LEV’s molecular mechanism of action remains unknown.

Consistent with its unique pharmacological profile, LEV has a novel target, Synaptic Vesicle protein 2A, (**SV2A**) (2). SV2A is one of three functionally redundant membrane proteins unique to secretory vesicles that undergo calcium-regulated secretion (3). SV2A is both necessary and sufficient for LEV binding (4), and multiple observations suggest that LEV acts by affecting SV2A function. This includes the fact that SV2A hypomorphic (SV2A^+/−^) mice are less responsive to LEV’s anti-seizure effects (5). Furthermore, the ability of structurally-related compounds to control seizures is directly correlated to affinity for SV2A (6). Although SV2 has been proposed to have multiple functions (7,8), its only fully vetted action is modulating the expression levels and trafficking of the calcium sensor protein synaptotagmin via a calcium- and phosphorylation-regulated interaction between the two proteins (9-11).

Of the three SV2 genes, SV2A is the most widely expressed. Notably, it is the only SV2 expressed in the majority of GABAergic neurons and dentate granule cells of the hippocampus (12,13), two classes of neurons implicated in the control of excitability. Loss of SV2A, but not other SV2 paralogs, results in severe seizures and early death (14,15). Interestingly, elevated SV2A is also associated with pathological neurotransmission. Seizure kindling in rodents is correlated with increased SV2A expression (16-18), and overexpression of SV2A in cultured neurons results in aberrant neurotransmission that is rescued by LEV (19). Thus, it appears that both decreased and increased SV2A may contribute to epilepsy.

LEV produces an activity-dependent decrease of both excitatory and inhibitory neurotransmission in hippocampal slice preparations (20,21). However, it has no effect on isolated, hippocampal neurons (19), suggesting that LEV action might not be via a direct effect on SV2A. Alternatively, because most neurons express both SV2A and SV2B, the lack of effect the presence of non-LEV binding SV2 paralogs may mask LEV’s effects. To begin to understand LEV’s mechanism of action at the cellular level, we examined LEV effects in neurons in which SV2A was the sole SV2 paralog. We report that in preparations from SV2B knockout (**BKO**) mice, LEV reduces synaptic depression in autaptic neurons, reduces SV2A binding to synaptotagmin, and slows synaptotagmin endocytosis in conventionally cultured hippocampal neurons. Our findings suggest that LEV acts primarily at synapses that express only SV2A by disrupting SV2A-regulated synaptotagmin function.

## Results

### LEV decreases synaptic depression in hippocampal neurons from BKO mice

Previous studies examining LEV effects on synaptic transmission in autaptic hippocampal neurons found no measureable effect on neurotransmission (19). The wild-type (**WT**) excitatory neurons used in those studies express SV2A and SV2B with low or non-existent levels of SV2C. Because all SV2 paralogs can rescue normal neurotransmission when expressed in neurons from SV2A^−/−^SV2B^−/−^ mice (**DKO**) mice (7), they are considered to be functionally redundant. Thus, the presence of SV2B, which does not bind LEV, might be masking measureable effects in WT excitatory neurons.

To assess LEV effects on neurotransmission in the absence of SV2B, we measured synaptic depression, an indicator of synaptic release probability (22). Excitatory principal neurons from the CA region of the hippocampus were cultured from SV2B^−/−^ (**BKO**) mice. Single neurons were grown on microislands of astrocytes allowing them to form autaptic synapses. Cultures were treated +/− 300uM LEV for 3-7 hours. This concentration and incubation time was chosen because it produced effects on synaptic transmission in hippocampal slices (23). Excitatory postsynaptic currents (EPSCs) in response to 2 sec, 20 Hz stimulus trains were recorded.

Cultured excitatory hippocampal neurons normally demonstrate synaptic depression during 20Hz stimulus trains. LEV treatment reduced the extent of this depression in neurons from BKO mice (**Figure 1A**). Consistent with this, the paired-pulse ratio, a comparison of the first two EPSCs, was increased by LEV treatment (PPR_Ctrl_ = 0.87±0.06, PPR_LEV_ = 1.02±0.04, *P <0.05) (**Figure 1B**). These changes in the rate of synaptic depression suggest that LEV is decreasing the probability of neurotransmitter release from BKO synapses (24). We note that this effect is similar to the decrease in synaptic depression observed in neurons lacking SV2 (25), suggesting that LEV impairs SV2 function. In contrast to its effects on neurons lacking SV2B, LEV had no significant effect on synaptic depression in neurons cultured from WT mice (PPR_ctrl_ = 0.82±0.05, PPR_LEV_ = 0.94±0.05) (**Figure 1C-D**). The selective effect of LEV on transmitter release from neurons expressing only SV2A suggests that the presence of other SV2 paralogs masks LEV effects in WT hippocampal neurons.

**Figure 1:**
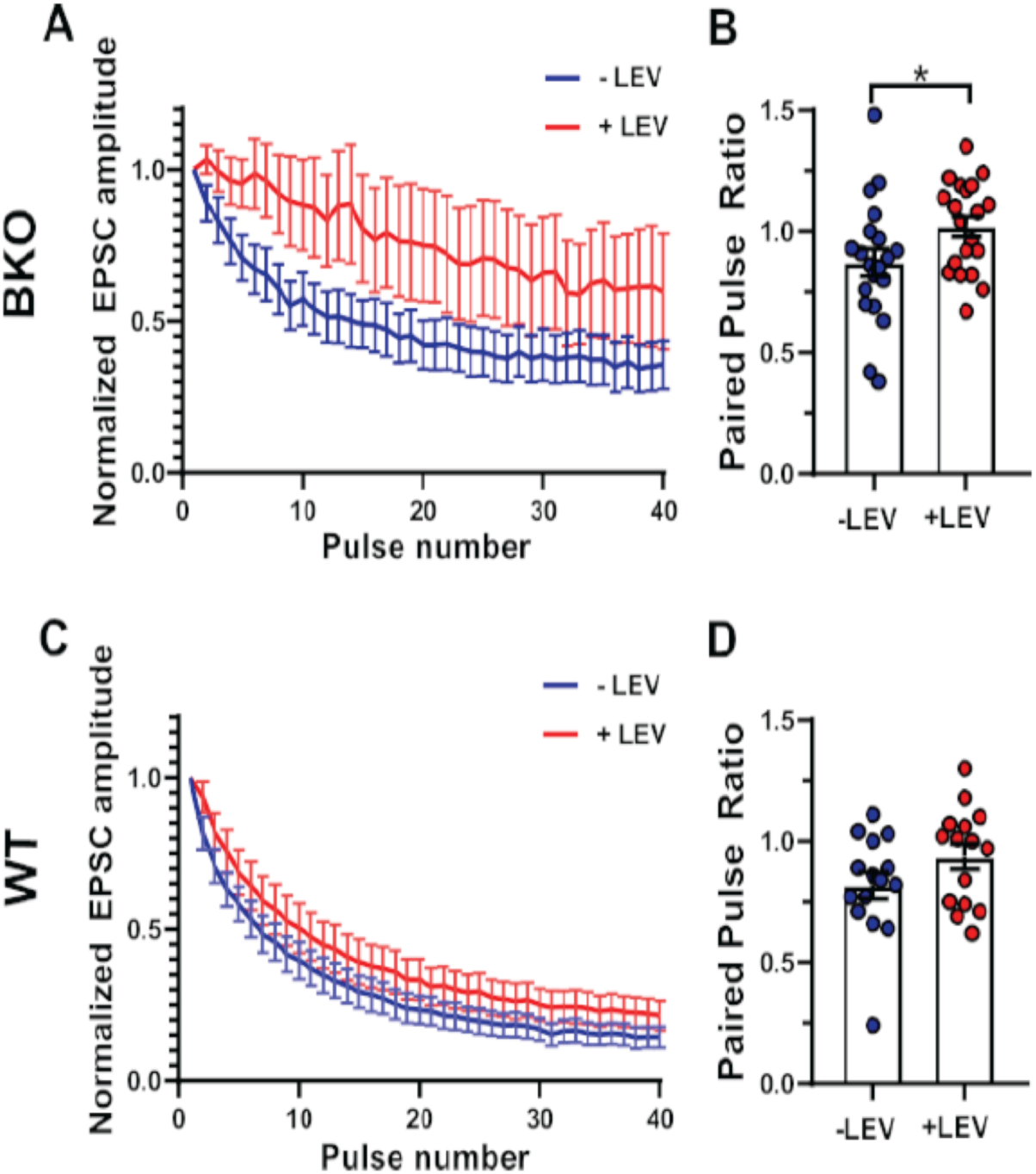
LEV decreases synaptic depression, an indicator of release probability, in isolated hippocampal neurons from SV2 BKO mice. Excitatory postsynaptic potentials (EPSCs) were measured using whole cell patch clamp stimulation and recording from isolated (autaptic) pyramidal neurons cultured on microislands of astrocytes. (A,C) Shown are averaged EPSC amplitudes in response to a 2s. 20Hz train. Amplitudes were normalized to the initial response then averaged. (A) BKO and (C) WT neurons treated +/− 300uM LEV for 3-7 hours. LEV decreased synaptic depression only in BKO neurons, suggesting that it reduces synaptic release probability in synapses that express only SV2A. (B,C) Paired pulse ratio, an index of synaptic release probability, was calculated as EPSC_2_/EPSC_1_. LEV (+LEV) increased the paired pulse ratio in neurons from BKO mice (PPR_+LEV_ = 1.02 ± 0.04, student’s t-test, \**P*<0.05, n = 20) compared to control (−LEV) (PPR_ctrl_=0.87 ± 0.06, n=20). In contrast, LEV had no effect on the paired pulse ratio in neurons from WT (PPR_+LEV_ ^=^ 0.94 ± 0.05, n=15, PPR_ctrl_ = 0.82 ± 0.05, n=15). Data represent the mean ± SEM.

### LEV decreases SV2A binding to synaptotagmin

SV2 binds the C2B domain of the calcium sensor protein synaptotagmin (10,26-29), domain essential to synaptotagmin’s role in exo (30-32) and endocytosis (33,34). To determine if LEV affects the interaction of SV2A with synaptotagmin, we assessed their binding *in vitro* in the presence and absence of 300uM LEV. Binding reactions included Triton X-100 extracts of mouse brain and resin coated with either recombinant glutathione-S-transferase (GST) or a glutathione-S-transferase–synaptotagmin 1 fusion protein (GST-Syt1)(27). Because calcium inhibits SV2 - synaptotagmin binding, we assayed LEV action in both low calcium (10^−9^M) and high (2mM) calcium. SV2A binding to the resin was then assessed by immunoblot analysis.

LEV treatment reduced SV2A binding to GST-Syt1 in both high and low calcium conditions when the source of SV2A was brain extracts of BKO mice. At low calcium concentrations, LEV reduced binding by ∼40%. Calcium alone decreased binding by 63%, and addition of LEV resulted in a further decrease to 14% of control (86% reduction) (**Figure 2A**). Somewhat surprisingly, LEV had no effect on SV2A binding to synaptotagmin when the source of SV2A was WT mouse brain (**Figure 2B**). The absence of LEV effects on SV2A when SV2B is present suggests that SV2s may form hetero-oligomers, precluding LEV action. Consistent with this, we find that SV2B co-immunoprecipitates with SV2A from extracts of WT mouse brain (data not shown).

**Figure 2:**
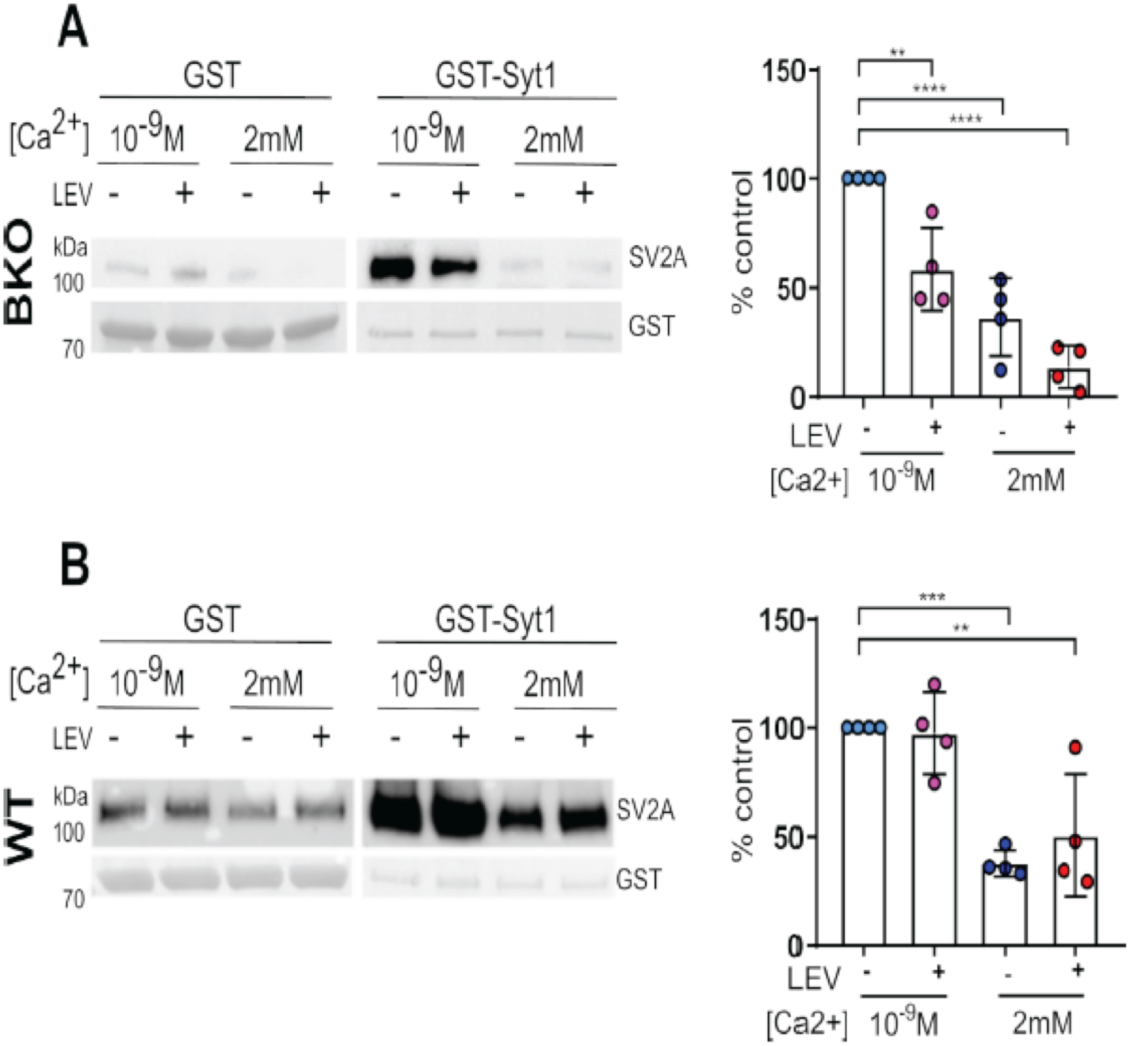
LEV decreases SV2A binding to synaptotagmin *in vitro* in the absence of SV2B. LEV effects of the SV2A-synaptotagmin interaction were assessed using a GST pull-down approach. The fusion protein GST-synaptotagmin 1 (GST-Syt1), or GST as a control, was attached to glutathione resin and incubated with extracts of mouse brain in the presence or absence of 300uM LEV and in low (10^−9^M) or high (2mM) calcium. SV2A binding was assessed by immunoblot analysis and normalized to the amount of GST-Syt1 or GST in the same lane. Binding was graphed as the percentage of the no LEV. low calcium (Control) condition. (**A**) SV2A from extracts of BKO mouse brain shows reduced binding to GST-Syt1 In the presence of LEV. Consistent with previous studies, high calcium reduced binding, and addition of LEV in the presence of high calcium further reduced binding (n=4, One-way ANOVA ** *P*<0.01,*** *P*<0.0001). Blots from a representative experiment are shown at the left. (**B**) SV2A from extracts of WT mouse brain showed reduced binding to GST-Syt1 in the presence of high calcium. In contrast to extracts from BKO mice. LEV had no effect on the interaction (n=4, One-way ANOVA \*\**P*<0.01,\*\*\**P*<0.0001,\*\*\*\**P*<0.0001). Blots from a representative experiment are shown at the left. Bar graphs represent data ± SD.

### LEV slows synaptotagmin endocytosis at synapses

The interaction of SV2A and synaptotagmin plays a crucial role in synaptotagmin’s ability to engage the proteins that mediate clathrin-dependent endocytosis (35). Thus SV2A is essential to synaptotagmin trafficking to synaptic vesicles (9,11). To determine if LEV’s inhibition of SV2A-synaptotagmin binding impacts synaptotagmin trafficking, we assessed its effects using two measures of synaptotagmin endocytosis in cultured hippocampal neurons.

First, we quantified the amount of surface stranded synaptotagmin in conventional hippocampal cultures treated with LEV. Cultures from WT or BKO mice were treated +/− 300uM LEV for 3 hours and then incubated with EZ-Link Sulfo-NHS-LC-LC-Biotin, a membrane-impermeant biotinylating reagent. After cultures were harvested and extracted in Triton X-100, surface stranded biotinylated proteins were isolated using streptavidin beads. The amount of biotinylated synaptotagmin and SV2A was quantified using Western blot analysis. Blots were probed for SV2A, synaptotagmin, and the plasma membrane protein sodium/potassium ATPase, which served as a loading control. Compared to untreated cells, LEV treatment increased the amount of synaptotagmin on the surface by ∼60%, suggesting decreased internalization (**Figure 3A**). Interestingly, there was no significant effect on the amount of biotinylated SV2A suggesting that SV2A’s internalization is not affected by LEV. Consistent with its effects on neurotransmission and *in vitro* protein interaction, LEV had no effect on the amount of biotinylated synaptotagmin or SV2A in WT neurons (**Figure 3B**). To control for the possibility that increased surface synaptotagmin was due to increased expression levels, we compared the amount of synaptotagmin in brain post nuclear supernatant from WT and BKO mice. There was no difference (data not shown). Thus, the most parsimonious interpretation is that LEV disrupts SV2A regulation of synaptotagmin trafficking.

**Figure 3:**
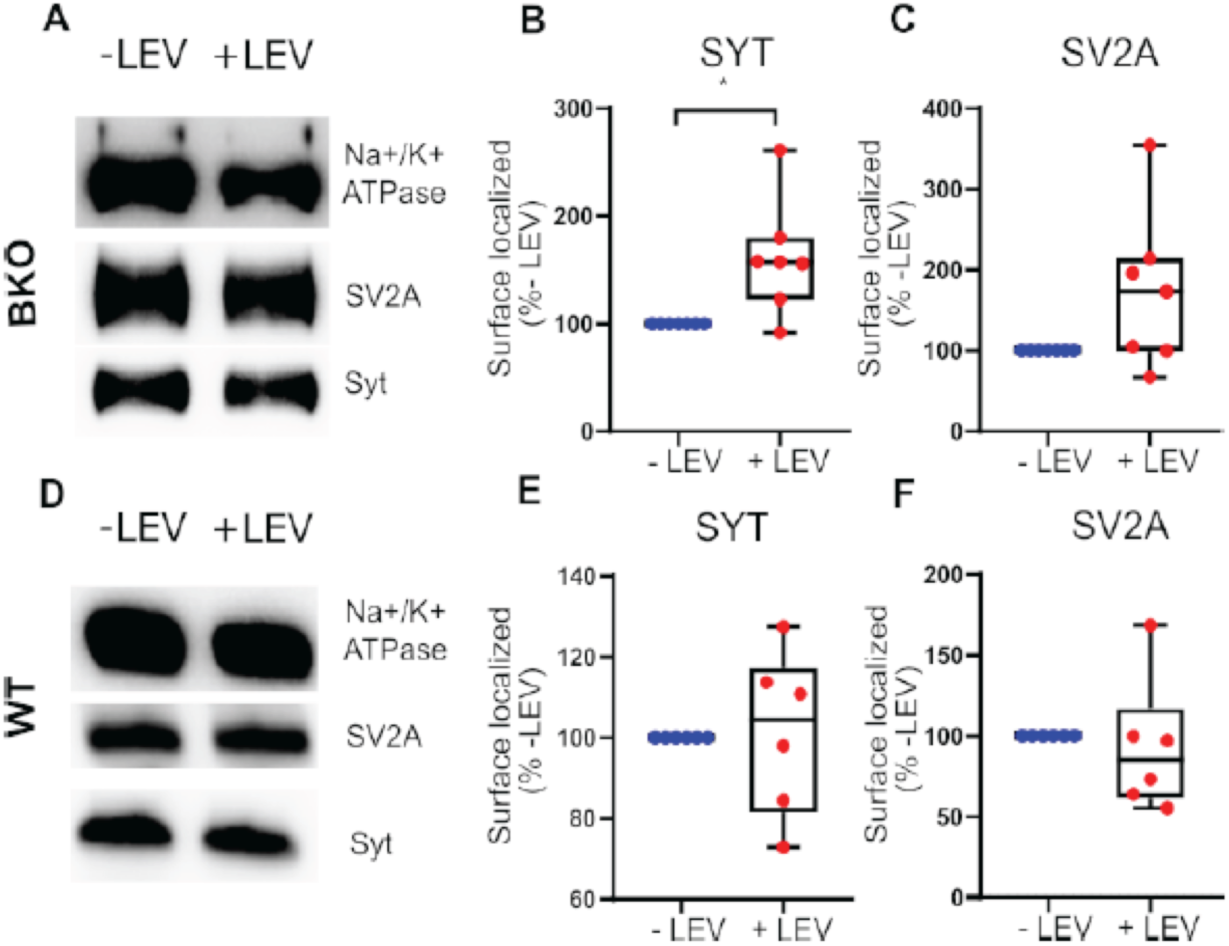
LEV increases surface localization of synaptotagmin in hippocampal neurons cultured from SV2 BKO mice. To assess LEV’s effect on SV2’s role in synaptotagmin trafficking, we compared the amount of surface localized synaptotagmin in hippocampal neurons treated +/-300uM LEV for three hours. Shown are representative immunoblots (A, D) and quantification of biotinylated synaptotagmin (Syt1) (B, E) and SV2A (C, F) in BKO and WT hippocampal neurons. To control for protein loading, the amount of biotinylated Syt1 or SV2A was normalized to biotinylated sodium potassium ATPase (Na+/K+ ATPase). Values were expressed as the percentage of the no LEV condition. (B) LEV increased the amount of Syt1 on the surface in neurons from BKO mice (Mann-Whitney, \**P*<0.05, n = 7) (C) LEV did not affect the amount of surface-localized SV2A. In neurons from WT mice, LEV had no significant effect on the amount of either Syt1 (E) or SV2A (F) in WT neurons (n=5). Each data point represents a different culture (n).

To measure LEV effects on the rate of synaptotagmin retrieval, we tracked the exo- and endocytosis of synaptotagmin1-pHluorin (Syt1-pHluorin), a fusion protein consisting of synaptotagmin 1 and pHluorin, a pH sensitive GFP variant (36). When fused to the lumenal domain of a synaptic vesicle protein, pHluorin is quenched by the low pH of the vesicle interior. It becomes fluorescent upon vesicle fusion and exposure to the neutral pH of the synapse exterior. As pHluorin fusion proteins are recycled back into vesicles during endocytosis, fluorescence is quenched. The rate of fluorescence decline can be used to measure the endocytotic time constant, tau.

Conventional hippocampal cultures from BKO mice were transfected with a construct driving the expression of Syt1-pHluorin under the control of the synapsin promoter. Prior to field stimulation, cultures were treated +/− 300uM LEV for 3-8 hours after which fluorescence was recorded at a rate of 2 Hz in response to a stimulus train of 200 action potentials delivered at 40Hz. LEV treatment slowed Syt1-pHluorin retrieval after the initial stimulus train. LEV increased the endocytic time constant by ∼60% (τ_ctrl_ = 53.9±5.4 s, τ_Lev_ = 86.7 ± 10.2 s) (**Figure 4A-C**). In contrast, LEV treatment had no significant effect on the time course of Syt1-pHluorin retrieval in cultures from WT mice (τ_ctrl_ = 49.69±5.18 s, τ_LEV_ = 55.6±7.44 s), consistent with it affecting only synapses that exclusively express SV2A (**Figure 4 D-E**).

**Figure 4:**
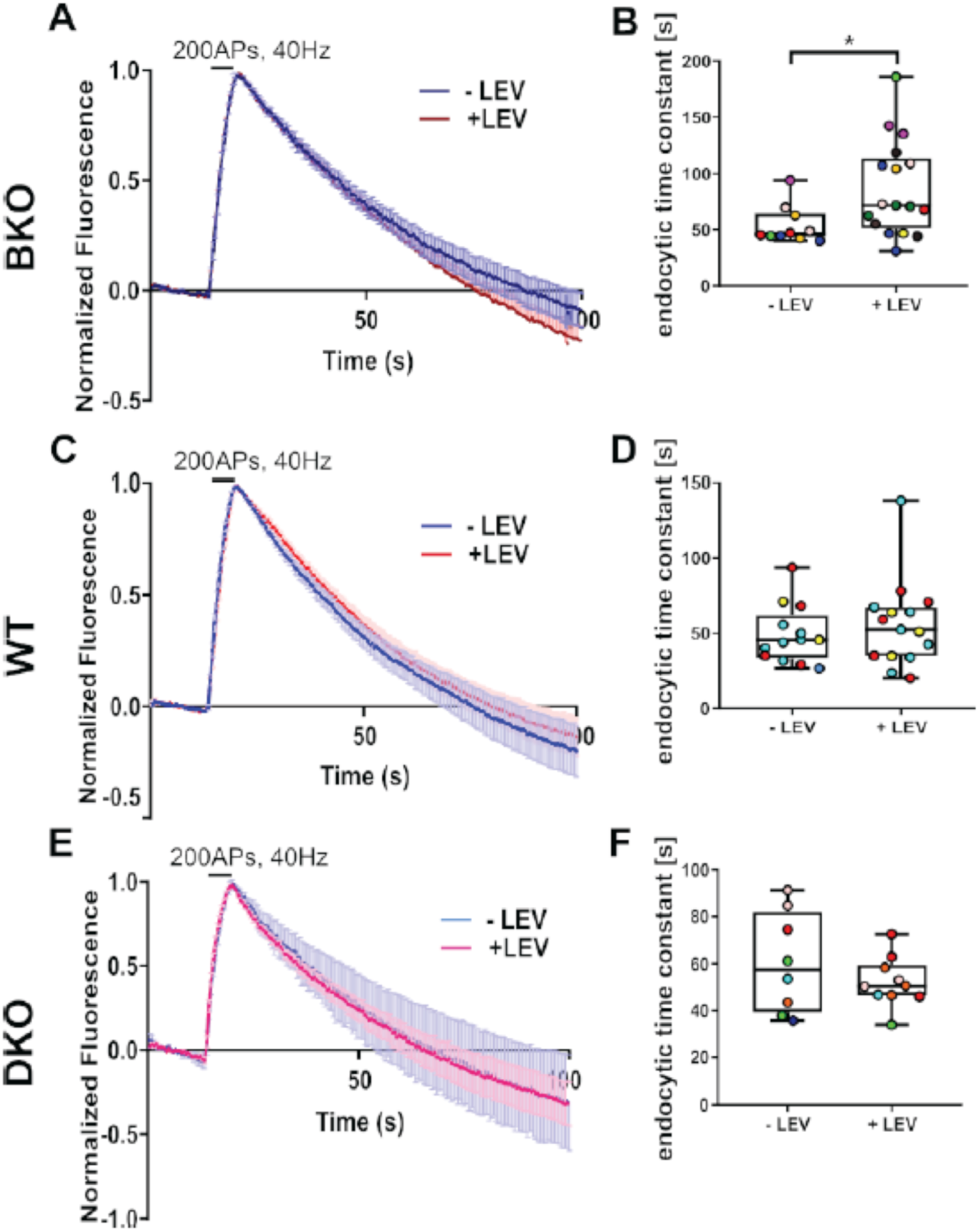
LEV slows Syt1-pHluorin retrieval in hippocampal neurons from SV2 BKO mice. To assess the effects of LEV on synaptotagmin trafficking, which is regulated by SV2A, we measured the fluorescence decay of Syt1-pHluorin in neurons from BKO, WT and DKO mice cultured on glass coverslips. Average time course (A, C, E) and endocytic time constants (B, D, F) of Syt1-pHluorin fluorescence in response to 200APs, delivered at 40Hz are shown. (A, B) In neurons from BKO mice, LEV increased the endocytic time constant (t) indicating slower endocytosis of Syt1-pHluorin (τ_ctrl_ = 53.9 ± 5.4 s, n = 6, τ_Lev_ = 86.7 ± 10.2 s, n = 9, Mann-Whitney, \**P* <0.05). (C, D) LEV had no effect on Syt1-pHluorin endocytosis in in WT neurons (τ_ctrl_ = 49.69± 5.18 s, n=4, τ_LEV_ = 55.6 ± 7.44 s, n=3).

Because complete loss of SV2A results in early death, it has been impossible to test the requirement for SV2A in LEV action. But because the cultures used for these studies are generated from neonatal mice (Postnatal days 0-2), we were able to compare LEV action in cultures from BKO from DKO mice. Consistent with a requirement for SV2A, LEV had no effect on the time course of Syt1-pHluorin retrieval in cultures from DKO mice (τ_ctrl_ = 60.2±7.55 s, τ_Lev_ = 52.13±3.34 s) (**Figure 4F-H**).

Loss of SV2A leads to a selective effect on the trafficking of synaptotagmin. To assess the specificity of the LEV action on endocytosis, we measured retrieval of synaptophysin, a vesicle protein whose trafficking is not linked to SV2 (9,11). For these studies, we tracked the exo- and endocytosis of synaptophysin-pHluorin (SypHy) in cultures from both WT and BKO mice. In contrast to LEV’s effects on Syt1-pHluorin trafficking, LEV had no significant effect on SypHy retrieval in either BKO (τ_ctrl_ = 61.17± 6.4 s, τ_LEV_ = 56.5±3.81 s) or WT (τ_ctrl_ = 55.45±6.92 s, τ_LEV_ = 45.92±3.92 s) cultures (**Figure 5**). This suggests that LEV has a selective effect on proteins that bind to SV2A.

**Figure 5:**
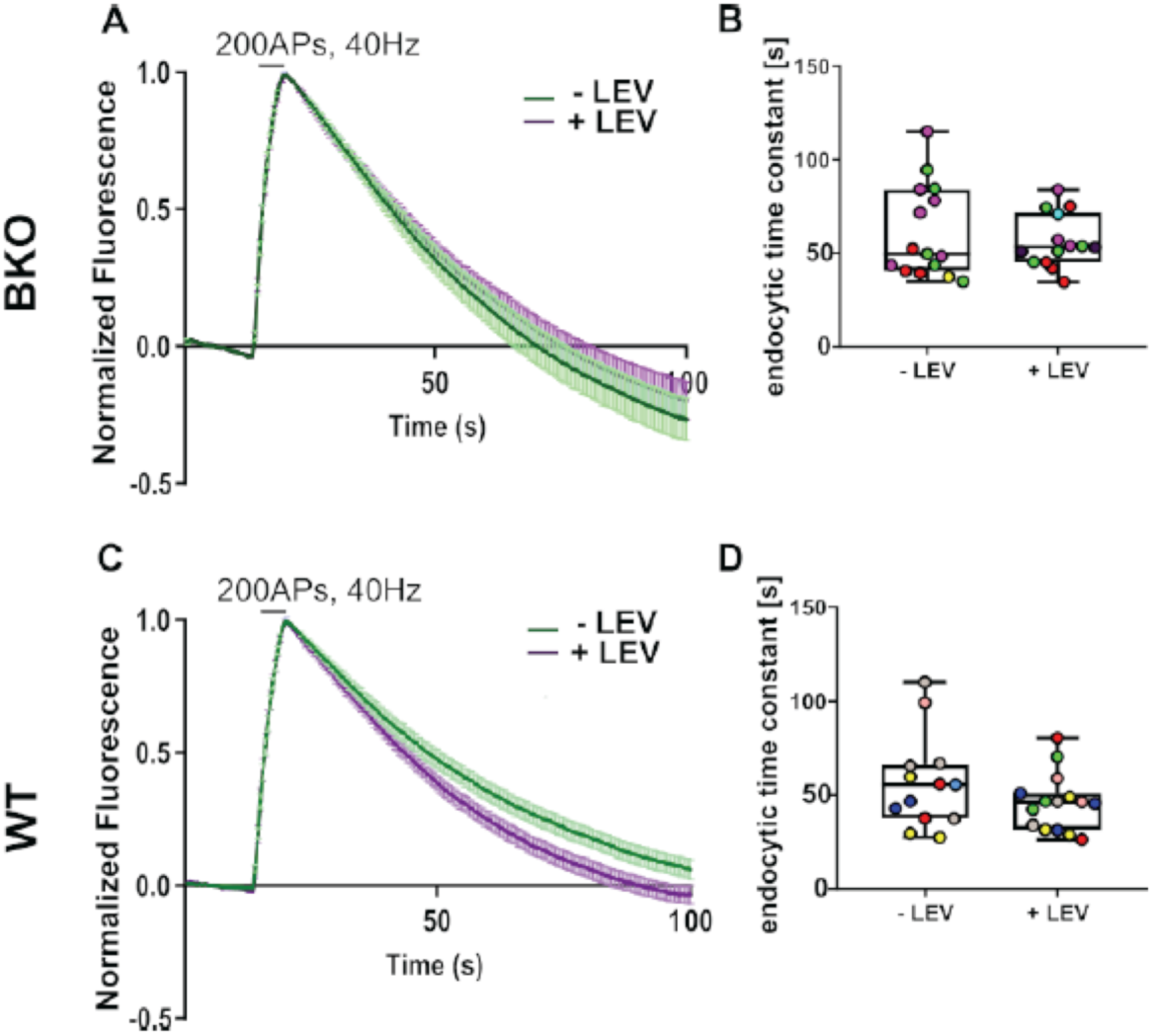
LEV does not affect SypHy retrieval in WT of BKO hippocampal neurons. To assess the effects of LEV on synaptophysin trafficking, which is not regulated by SV2A, we measured the fluorescence decay of synaptophysin pHluorin (SypHy) in neurons from BKO, WT and DKO mice cultured on glass coverslips. Average time course (A, C) and endocytic time constants (B, D) of SypHy fluorescence in response to 200APs, delivered at 40Hz are shown. Average time course (A,C) and endocytic time constants (B,D) of synaptophysin (SypHy) in BKO (A,B) and WT (C,D) hippocampal neurons in response to 200APs, 40Hz. LEV did not affect SypHy endocytosis in BKO neurons (τ_ctrl_ = 61.17 ± 6.4 s, n = 4,τ_LEV_ = 56.5 ± 3.81 s, n = 5) or in WT neurons (τ_ctrl_ = 55.45 ± 6.92 s, n = 2, τ_LEV_ = 45.92 ± 3.92 s, n = 3). Average time course are graphed as mean ± SEM. Endocytic time constants are graphed using box plots. Culture dates (n) are indicated by color. Each data point represents a separate coverslip.

## Discussion

The work presented here provides two insights into the mechanism LEV action. First, we show that LEV action is affected by the presence of other SV2 paralogs. suggesting that LEV action predominates at synapses that express SV2A exclusively. This finding has important implications not only for understanding how LEV reduces seizure activity, but also for the development of compounds that target other SV2 paralogs. Second, we demonstrate that LEV acts at the molecular level by disrupting SV2A’s interaction with the calcium sensor protein synaptotagmin. These findings provide further evidence that LEV action is unique among anti-epileptic medications.

### LEV action predominates at synapses that express only SV2A

A standing puzzle in the search for LEV’s cellular action has been the absence of an effect on isolated neurons (19), which precluded genetic or structure-function studies of LEV action at the cellular level. Our finding that LEV reduced synaptic depression in isolated hippocampal neurons from BKO but not WT mice allows us to re-evaluate previous hypotheses of LEV action.

Selectivity for synapses that contain only SV2A suggests that LEV acts narrowly in the brain, despite the widespread expression of SV2A. SV2A is co-expressed with SV2B in most brain synapses. The exceptions are the majority of GABAergic synapses and the excitatory projections of dentate granule neurons in the hippocampus, which contain only SV2A (12,13). Both types of synapses have been implicated in the control of brain excitability (37,38). The targeted action of LEV at synapses that play a major role in the development and control of seizures may be the basis for its efficacy and high therapeutic index. Furthermore, because SV2A expression has been reported to increase in some forms of epilepsy (18,39), determining whether LEV has more widespread action in epileptic brain will be important to refining its therapeutic use.

Because our preparation utilizes neurons before the early death induced by loss of SV2A, we were able to rigorously test the requirement for SV2A in LEV action. The absence of a LEV effect in neurons from DKO mice is the first demonstration that loss of SV2A abolishes LEV action.

### LEV inhibits SV2A binding to synaptotagmin

Although multiple functions have been ascribed to SV2 (8,40-42), it has one verified function. SV2 binds to and regulates the stability and trafficking of synaptotagmin (9,11,26,29,43). The fact that synaptotagmin expression levels and trafficking are disrupted with loss of SV2 suggests that SV2 plays an escort or chaperone-like function. Because synaptotagmin plays an essential role in both exo- and endocytosis, it suggests that LEV’s ultimate mechanism of action may be the disruption of synaptotagmin function. This interpretation is consistent with the previous finding that LEV restores normal synaptotagmin levels in synapses overexpressing SV2A (19). It also supports the conclusion that regulation of synaptotagmin is a primary function of SV2 (44). Thus, LEV-mediated inhibition of the SV2A-synaptotagmin interaction could have a direct effect on synaptotagmin’s ability to engage with the vesicle fusion (SNARE) complex (32).

Synaptotagmin plays an essential role in regulating vesicle exocytosis by interacting with membrane lipids and the proteins that mediate vesicle fusion (SNAREs) (32). By disrupting SV2A’s chaperone or escort action, LEV could affect synaptotagmin’s ability to contribute to exocytosis. Consistent with this, we note that loss of SV2, which LEV treatment recapitulated in our studies of synaptic plasticity, reduces synaptic vesicle priming and results in fewer assembled vesicle fusion (SNARE) complexes (25,45).

Synaptotagmin has also been implicated in endocytosis (34). SV2A promotes the binding of synaptotagmin to AP2, a component of the clathrin-dependent endocytosis machinery (35). Our observation that LEV slows synaptotagmin endocytosis and decreases its internalization suggests that by disrupting the SV2A–synaptotagmin interaction, LEV’s mechanism of action includes affecting synaptotagmin retrieval. This suggests that LEV could act by decreasing the amount of synaptotagmin in recycling vesicles. The interpretation that LEV reduces the replenishment of release competent vesicles is consistent with previous studies reporting that LEV produces an activity-dependent decrease in synaptic transmission (20) via faster supply rate depression (46). In contrast, a LEV effect on synaptotagmin recycling or function suggests an alternate explanation for the observation that LEV requires exocytosis in order to be effective. Rather than requiring vesicle exocytosis to get LEV into synapses (21), our finding that LEV disrupts SV2A binding to synaptotagmin suggests that LEV action is limited to vesicles that have yet to be formed and/or primed for release.

In summary, our results provide initial insights into the molecular action of LEV. Further analysis of how LEV impacts protein interactions in presynaptic terminals will be the next stage towards developing a structural approach to further drug development.

## Experimental Procedures

### Animals and reagents

The generation of SV2A^+/+^SV2B^−/−^ (BKO) and SV2A^−/−^SV2B^−/−^ (DKO) mice was reported previously (25). Wild-type (WT) and BKO mice were kept as separate colonies generated from littermate heterozygous crosses. All animals were on a 99.99% C57BL/6 genetic background. DKO mice were generated by crossing SV2A^+/−^SV2B^−/−^ mice. Genotype was determined by PCR before culturing neurons. The animal protocol was reviewed and approved by the Institutional Animal Care and Use Committee of the University of Washington.

Levetiracetam (LEV) was from Sigma-Aldrich (L8668-100MG). Polyclonal anti-SV2A antibody was house generated and affinity purified from sera (13), rabbit polyclonal anti-synaptotagmin antibody was raised against the cytoplasmic domain, and anti-Na+/K+ ATPase was from Development Studies Hybridoma Bank (a6F). Antibody binding in immunoblot analyses was detected using horse radish peroxidase-conjugated secondary antibodies and visualized using enhanced chemiluminescent chemistry (BioRad Clarity™) and visualized via Bio-Rad Chemidoc. Immunoreactive bands were quantified using ImageLab software (BioRad). Secondary antibodies were used at a 1: 5000 dilution (Invitrogen HRP-conjugated goat anti-rabbit G-21234 and HRP-conjugated goat anti-mouse G-21040).

### Tissue Culture/Transfections

For electrophysiological recordings, neurons isolated from the CA region of the hippocampus of P0-P1 mice were cultured on small microislands of permissive substrate as previously described (25). Neurons were plated on a feeder layer of astrocytes, then grown in neural medium without mitotic inhibitors and used for recordings after 12-16 days in vitro (DIV). For all other experiments, CA hippocampal neurons were cultured on 25mm coverslips coated with poly-D-lysine and rat collagen and layer of astrocytes as previously described (9).

Cultures were transfected using calcium phosphate at 6-8 DIV. Synaptotagmin1-pHluorin (Syt1-pHluorin) was obtained from Dr. Volker Haucke (Leibniz Institute of Molecular Pharmacology). Synaptophysin-pHluorin (SypHy) was obtained from Dr. Leon Lagnado (University of Sussex). For transfections, cultures were serum starved for 1.5 hours prior to transfection. A calcium phosphate/DNA precipitate was formed in HEPES-buffered saline [280mM NaCl, 10mM KCl, 1.5mM NaH_2_PO_4_, 12mM D-glucose, 50mM HEPES, pH 7.08] for 20-25 minutes. The precipitate (200uL) was added drop-wise to 2mL serum free media and incubated for 1.5 hrs. After transfection, the cultures were returned to their original culture medium.

### Electrophysiology

Following treatment +/− 300 uM LEV for 3-7 hrs, whole-cell voltage clamp recordings were made from excitatory neurons on single-neuron islands using an Axopatch 200A amplifier (Axon Instruments, Sunnyvale, CA). The extracellular recording solution contained 119 mM NaCl, 5 mM KCl, 2 mM CaCl_2_, 1.5 mM MgCl_2_, 30 mM glucose, 20 mM HEPES, and 1 uM glycine; 300 uM LEV or water (vehicle) was added to the external solution of cells during recording. The pipette solution contained 148.5 mM K-gluconate, 9 mM NaCl, 1 mM MgCl_2_, 10 mM HEPES, and 0.2 mM EGTA. Cells were held at −60 mV and stimulated with a 1 ms depolarization to +20 mV to evoke neurotransmitter release. Recording electrodes were 2.5-3.5 MΩ, and series resistance was compensated 70-85%. Cells with series resistance > 20 MΩ before series resistance compensation, or leak current >250 pA were excluded. All experiments were performed at room temperature.

### Glutathione S-transferase (GST) pull-down assays

Protein binding (pull-down) assays were performed as previously described (27). Brains from WT or BKO mice were homogenized in HBS buffer [10mM HEPES, 142mM NaCl, 2.4mM KCl, 1mM MgCl_2_, 5mM D-glucose, 1mM EGTA, 1x protease inhibitor (Roche)]. The brain homogenate was centrifuged at 1000xg for 10 minutes to remove nuclei and undisrupted cells. One milligram of the resulting post nuclear supernatant was solubilized in an equal volume of solubilization buffer [20mM HEPES, 1mM EGTA, 95mM potassium acetate, 2% Triton X-100, with or without 3mM CaCl2 and 300µM LEV or water] for two hours on ice then centrifuged at 19,000xg for 30 minutes at 4°C. The resulting supernatant (extract) was incubated with bacterially-produced GST-synaptotagmin1 (GST-Syt1) bound to Pierce™ glutathione agarose (16100) for 1 hour at 4°C. After binding, beads were washed four times with 1mL wash buffer [10mM HEPES, 1mM EGTA, 47.5mM potassium acetate, 1% Triton X-100, with or without 3mM CaCl2 and 300µM LEV or water]. Proteins were eluted with 8M urea dissolved in 100mM Tris-HCl, pH 8, or SDS-PAGE sample buffer, and subjected to Western analysis.

### Surface Biotinylation

Hippocampal cultures (14-16 DIV) from WT or BKO mice were treated with 300uM LEV or water in neural media at 37°C for three hours. After incubation, cells were placed on ice and washed with cold phosphate buffered saline (PBS) [137mM NaCl, 2.7mM KCl, 10mM NaH_2_PO_4_, 1.8mM KH_2_PO_4_]. To biotinylate surface stranded cells, cells were incubated with 2mM EZ-LINK Sulfo-NHS-LC-LC-Biotin (Thermo Scientific 21338) for 25 minutes at 4°C. Cells were washed with 100mM glycine in PBS for 15 min to quench the reaction and then washed with PBS twice. Cells were harvested and extracted in lysis buffer [10mM HEPES, 140mM NaCl, 1% Triton X-100 and protease inhibitor] for 30 minutes at 4°C and spun at 19,000xg for 20 minutes at 4°C. Biotinylated proteins were isolated using Streptavidin Sepharose ® High Performance resin (GE 17-5113-01) and eluted with SDS-PAGE sample buffer. Eluted proteins were then subjected to immunoblot analysis.

### Imaging

Imaging of live cultures was performed at 14-16 DIV. Prior to imaging, neurons were incubated in neural medium with 300uM LEV or water for 3-8 hours at 37°C. Cultures were perfused in imaging buffer [170mM NaCl, 3.5mM KCl, 0.4mM KH_2_PO_4_, 20mM N-Tris(hydroxy-methyl)-methyl-2-aminoethane-sulphonic acid (TES), 5mM NaHCO_3_, 5mM glucose, 1.2mM Na_2_SO_4_, 1.2mM MgCl_2_, 1.3mM CaCl_2_, 10µM CNQX, and 50µM AP-5, pH 7.4] +/− 300uM LEV prior to imaging. To evoke synaptic transmission, neurons were subjected to electrical field stimulation using an RC21-BRFS chamber (Warner Instruments). A train of 200 action potentials was delivered at 40Hz using a Grass SD9E Stimulator. Fluorescent images (100ms exposures) were captured using an Olympus IX-10 microscope equipped with a 60x water-immersion objective and an Andor scMOS camera. Images were acquired at 2 Hz using Micro-Manager software and analyzed using NIH ImageJ (with Time Series Analyzer V3 plugin). Responding boutons were identified and total fluorescence intensity monitored over time. Fluorescence intensity was corrected for background (47). Endocytic time constants were determined by fitting the mono-exponential decay of fluorescence intensity using Prism (Graphpad) software.

### Statistical Methods

All statistical analyses were performed using Prism (Graphpad) software. Data analysis was performed with the assistance from the Department of Biostatistics Consulting Service at the University of Washington.

## Acknowledgements

This work was supported, in part, by grants from the National Institutes of Mental Health (R21MH099741 to SB), National Institutes of General Medical Sciences (T32GM007270 and T32GM007750 to KC).

